# JDINAC: joint density-based non-parametric differential interaction network analysis and classification using high-dimensional sparse omics data

**DOI:** 10.1101/099234

**Authors:** Jiadong Ji, Di He, Yang Feng, Yong He, Fuzhong Xue, Lei Xie

## Abstract

**Motivation:** A complex disease is usually driven by a number of genes interwoven into networks, rather than a single gene product. Network comparison or differential network analysis has become an important means of revealing the underlying mechanism of pathogenesis and identifying clinical biomarkers for disease classification. Most studies, however, are limited to network correlations that mainly capture the linear relationship among genes, or rely on the assumption of a parametric probability distribution of gene measurements. They are restrictive in real application.

**Results:** We propose a new Joint density based non-parametric Differential Interaction Network Analysis and Classification (JDINAC) method to identify differential interaction patterns of network activation between two groups. At the same time, JDINAC uses the network biomarkers to build a classification model. The novelty of JDINAC lies in its potential to capture non-linear relations between molecular interactions using high-dimensional sparse data as well as to adjust confounding factors, without the need of the assumption of a parametric probability distribution of gene measurements. Simulation studies demonstrate that JDINAC provides more accurate differential network estimation and lower classification error than that achieved by other state-of-the-art methods. We apply JDINAC to a Breast Invasive Carcinoma dataset, which includes 114 patients who have both tumor and matched normal samples. The hub genes and differential interaction patterns identified were consistent with existing experimental studies. Furthermore, JDINAC discriminated the tumor and normal sample with high accuracy by virtue of the identified biomarkers. JDINAC provides a general framework for feature selection and classification using high-dimensional sparse omics data.

**Availability:** R scripts available at https://github.com/jijiadong/JDINAC

**Supplementary information:** Supplementary data are available at bioRxiv online.

## 1 Introduction

It is well known that a complex biological process, such as the development and progression of cancer, is seldom attributed to a single molecule. Numerous cellular constituents, such as proteins, DNA, RNA and small molecules do not function in isolation, but rather interact with one another to fulfill particular biological functionality. In the view of network biology (Yoshimura *et al.*, 1998; Zhou *et al.*, 2011), a cellular function is a contextual attribute of quantifiable patterns of interactions between myriad of cellular constituents. Such interactions are not static processes, instead they are dynamic in response to changing genetic, epigenetic, and environmental factors (Bandyopadhyay *et al.*, 2010; Califano, 2011). The molecular interactions can be effectively abstracted as a network. Thus, differential network analysis becomes an important tool to understand the roles of different modules in complex biological processes, and draws tremendous attention. Typically, differential genetic interactions are a reflection of which cellular processes are differentially important under the studied condition (de la Fuente A, 2010; Ideker and Krogan, 2012).

In the past decade, many methods have been proposed to detect the differential network connection patterns between two condition-specific groups (e.g. patients and health controls). Gambardella *et al.* (2013) introduced DINA procedure to identify whether a known pathway is differentially co-regulated between different conditions. Yates and Mukhopadhyay *et al.* (2013) provided a dissimilarity measure that incorporates nearby neighborhood information for biological network hypothesis tests. Recently, Ruan *et al.* (2015) developed the dGHD algorithm for detecting differential interaction patterns in two-network comparisons. All of the aforementioned methods endeavor to identify whether the global network topology changed significantly between two groups. However, it will be of benefit to reveal critical pairwise molecular or genetic interactions that are responsible for the different physiological or pathological states of an organism in many applications. The identification of such interactions may help us to illuminate the underlying genetic mechanisms of complex diseases (e.g. cancer), to predict drug off-target effects (T Evangelidis, 2014), to develop multitarget anti-cancer therapy (Xie and Bourne, 2015), and to discover clinical biomarkers for disease classification.

To this end, the primary focus in this paper is to identify pairwise differential interactions among genes that are most closely related to a certain disease status. Most of such studies first require to divide the data into two separate groups according to the factor of interest. Besides, a certain correlation metric is often involved to represent the strength of pairwise interaction between nodes in the network. The existing methods mainly fall into two categories. The first category is to compare topological characteristics, such as degree, clustering coefficient of vertices within the network, of the constructed sparse network on grouping specific data (Reverter *et al.*, 2006; Zhang *et al.*, 2009). The main challenge of this approach lies in how to select appropriate threshold for constructing sparse network, although there have been miscellaneous methods proposed to address this challenge (Carter *et al.*, 2004; Elo *et al.*, 2007). To the best of our knowledge, no commonly feasible approach has been available yet. Approaches in the second category normally handle weighted group-specific network to further construct the differential network. In one manner such approach can only concentrate on edge-level to construct edge-difference based differential network (Hudson *et al.*, 2009; Liu *et al.*, 2010; Tesson *et al.*, 2010). On the other hand, it could focus more on finding gene sets and identify correlation patterns difference between groups. For example, the CoXpress (Watson, 2006) first performs hierarchical clustering with correlation matrix obtained from normal samples (or disease sample), then applies statistical test to determine whether the average correlation within one cluster is higher (or lower) than expected by chance and thus finally identifies the differentially co-expressed gene groups. Similarly, DiffCorr (Fukushima, 2013) identifies the first principle component based ‘eigen-molecules’ in the correlation matrices constructed from the grouped dataset, then performs Fisher z-test between the two groups to discover differential correlation. In addition, Zhao *et al*.(2014) proposed a direct estimation method (DEDN), which modeling each condition-specific network using the precision matrix under Gaussian assumption. However, most of the methods mentioned above are based on marginal or partial correlation. It can only capture the linear relationship among genes, which could be restrictive in real applications. It is often the case that nonlinear relationships exist between genes. Another critical but inadequately addressed issue is how to adjust the confounding factors in the differential network analysis. For instance, the condition-specific label is the length of the survival time of cancer patients, one group are patients with longer survival time and the other group are those with short survival time. Then the age of the patients is a potential confounding factor which needs to be adjusted. If the patients’ ages are different between two groups, it’s hard to know whether the identified differential network is associated with the survival time or the age. Furthermore, how to use the identified network biomarkers to achieve classification still poses great challenge in discriminant analysis especially in high-dimensional settings (He *et al,* 2016).

To address the challenges in differential network analysis and classification mentioned above using high dimensional sparse omics data, we propose a Joint density based non-parametric Differential Interaction Network Analysis and Classification (JDINAC) method to identify differential patterns of network activation between condition-specific groups (e.g. patients and health controls). The contribution of our work lies in that we can not only deal with the nonlinear relationship between the genes but also adjust the confounding factors in the differential network analysis. Furthermore, JDINAC is free of the assumption of a parametric probability distribution of gene measurements. We compare the ability of identifying differential network of our methods with DiffCorr (Fukushima, 2013), DEDN (Zhao *et al.*, 2014) and Lasso based method. By integrating the logistic regression into our method, our method is capable of accurate classification using high-dimensional sparse data. We also compared the classification performance of our method with Random Forest (RF) (Breiman, 2001), Naive Bayes (NB) and Lasso based methods in both simulation studies and real data example.

## 2 Methods

Network differential analysis and classification using high-dimensional sparse omics data face several challenges. Firstly, the number of data points *n* is often much smaller than the number of features *p*, e.g. *p*>>*n* problem. Secondly, the relationship between two biological variables is often non-linear. Thirdly, confounding factors often need to be adjusted in the differential network analysis and classification. Finally, the underlying distribution of biological variable may not follow Gaussian or other probability distribution on which many algorithms are based. JDINAC is proposed to address these problems.

Assume that we have observed gene-level activities (such as mRNA, methylation or copy number) for *p* genes measured over individuals. For individual *l* (*l* = *1, 2*, …, *N*), the binary response variable is denoted as
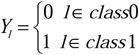 and the expression level of *i*^th^ gene is denoted as *x_li_*.

The JDINAC approach based on the logistic regression model can be constructed as,

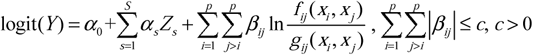

where *Z_s_* (*s* = 1,^…^, *S*) denote the covariates (e.g. age and gender). *f_ij_* and *g_ij_* denote the class conditional joint density of *x_i_* and *x_j_* respectively for class 1 and class 0, i.e., ((*x_i_*, *x_j_*)|*Y* = 1) ~ *f_ij_* and ((*x_i_*, *x_j_*)|*Y* = 0) ~ *g_ij_*. The conditional joint densities *f_ij_* (*x_i_*, *x_j_*) can indicate the strength of association between *x_i_* and *x_j_* in class 1. Since the number of pairs (*x_i_*, *x_j_*) can be larger than the sample size, the *L*_1_ penalty (Tibshirani, 1996) was adopted in this high-dimensional setting. Note that the above formulation can be viewed as an extension of the FANS approach (Fan *et al.*, 2016). Parameters *β_ij_* ≠ 0 indicate differential dependency patterns between condition-specific groups.

*L*_1_ regularized estimate for ***β***:

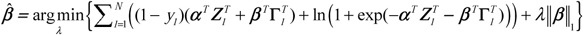

where

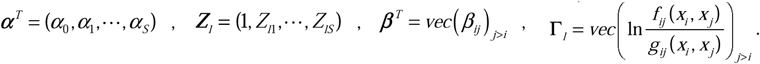

The advantages of the JDINAC approach over existing methods lie in the following aspects:1) it can achieve differential network analysis and classification simultaneously;2) it can adjust confounding factors in the differential network analysis, for example, if the samples are from cancer patients with different length of survival time, then the age of the patient is a potential confounding factor which needs to be adjusted.3) it is a nonparametric approach and can identify the nonlinear relationship among variables. Besides, it does not require any conditions on the distribution of the data, which makes it more robust.

JDINAC can be implemented as follows with its workflow shown in Figure 1.

**Fig 1.**
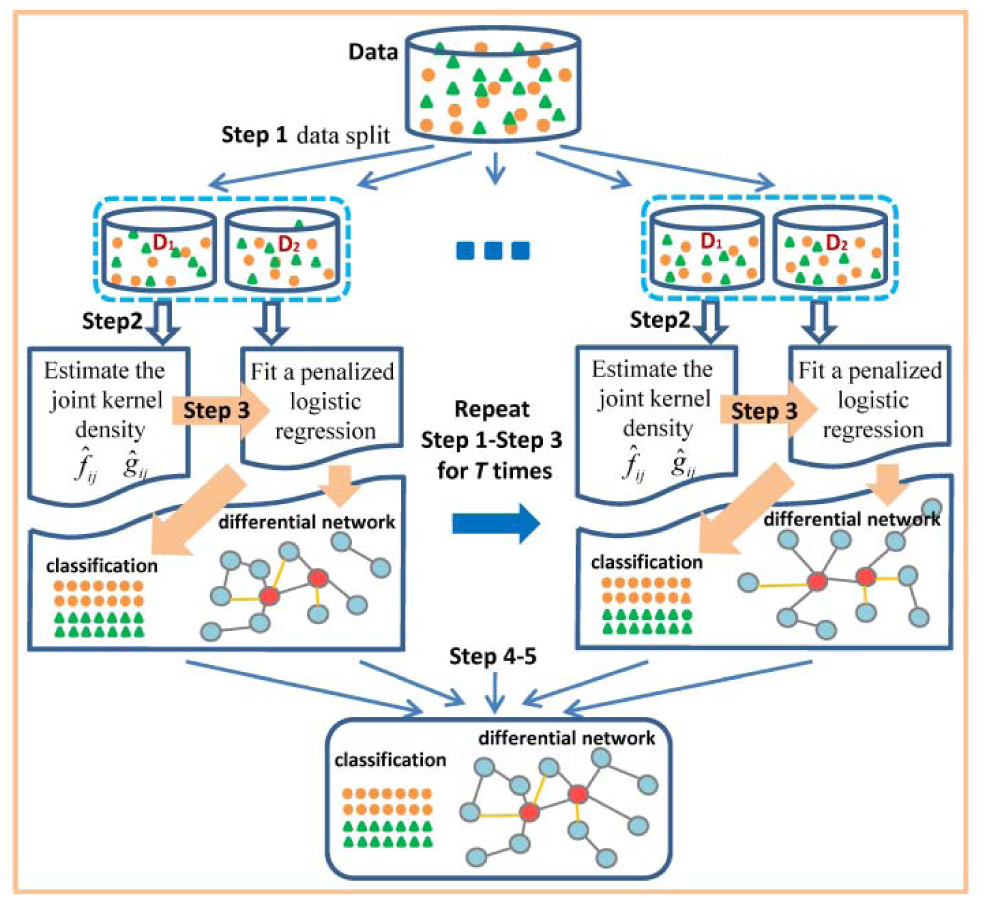
Workflow of JDINAC. Step 1. Given *N* observations *D* = {(*Y_l_*, ***X**_l_*), *l* = 1,…, *N*}. Randomly split the data into two parts: *D* = (*D*_1_, *D*_2_). Step 2. On part *D*_1_, estimate the joint kernel density functions 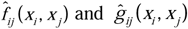, *i*,*j* = 1,…, *p*, *j* > *i*. Step 3. On part *D*_2_, fit an *L*_1_ -penalized logistic regression 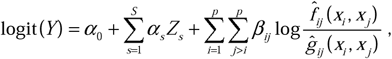 using cross validation to get the best penalty parameter. Step 4. Repeat Step 1-Step 3 for *T* times, for individual *l* using the average prediction 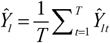 as the final prediction, and assign the *l*^th^ individual to class 1 if *Ŷ_l_* > 0.5, and class 0 otherwise. Setp 5. Calculate the differential dependency weight of each pair (*x_i_*, *x_j_*,) between two groups, 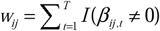, *i*, *j* = 1,…, *p*, *j* > *i*; where *I*() is the indicator function.

### 2.1 Simulation studies

Four simulation scenarios were designed for assessing the performances of differential network analysis and classification accuracy. In scenarios 1 and 2, the difference of association strength between pairs of genes in a network is caused by the different correlation (Figure 2a, Figure 2b). In scenario 3, the differential pairs have the same correlation structure between condition-specific groups but different joint density (Figure 2c). In scenario 4, the differential strength of association between pairs of genes in a network is caused by the nonlinear dependence (Figure2d). For scenario 1, 2 and 3, we generated 100 pairs of datasets, each representing the case (class 1) and the control (class 0) conditions. Each dataset contains 300 observations with *p* variables drawn from the multivariate normal distribution with mean 0 and covariance matrix Σ, that is, *X* ~ *N_p_*(**0**, Σ). Σ consists of 3 blocks along the diagonal. Σ = *diag*(Σ_1_, Σ_2_, Σ_3_), Σ_1_ = (*σ_ij_*)_*m*×*m*_, *σ_ij_* = *ρ*^|*i*–*j*|^ for *i*, *j* = 1, …, *m*; *m* = 80; 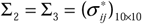.

**Fig 2.**
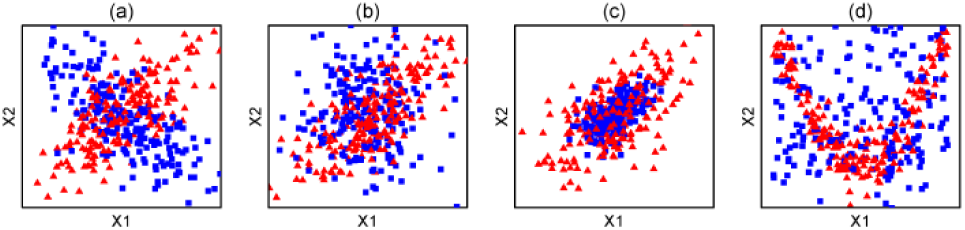
The scenarios of simulation studies. The blue square and red triangle represents the scatter plots for the two variables in class 0 and class 1 respectively, (a) scenario 1, the two variables is negatively correlated in class 0 and positively correlated in class 1, (b) scenario 2, the two variables are correlated in one group and are independent in the other, (c) scenario 3, the two variables are equally correlated but with different density in the two groups, (d) scenario 4, the two variables are independent in one group and have nonlinear relationship in the other group.

Scenario 1: In class 0, *p* = 100, *ρ* = 0.5, 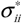 = 1 for *i* = 1,…,10, 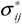 = (−1)^|*i*, *j*|^ × 0.5 for *i* ≠ *j*; in class 1, *p* = 100, *ρ* = 0.5, 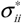 = 1 for *i* = 1, …, 10, 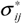 = 0.5 for *i* ≠ *j*.

Scenario 2: In class 0, *p* = 100, *ρ* = 0.5 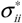 = 1 for *i* = 1,…, 10, 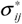 = 0 for *i* ≠ *j*; in class 1, *p* = 100, *ρ* = 0.5, 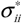 = 1 for *i* = 1,…, 10, 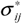 = 0.7 for *i* ≠ *j*.

Scenario 3: In class 0, *p* = 100, *ρ* = 0.5, 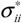 = 1 for *i* = 1,…, 10, 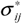 = 0.6 for *i* ≠ *j*; in class 1, *p* = 100, *ρ* = 0.5, 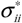 = 5 for *i* = 1, …, 10, 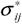 = 3 for *i* ≠ *j*.

Scenario 4: In class 0, generate data 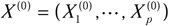, where 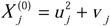, *j* = 1, …, *p*/2, *U_j_* ~ *Unif*(−2, 2) and *v_j_* ~ *N*(−4/3, 1/4); 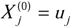, *j* = *p*/2+1,…, *p*, *p* = 100. In class 1, generate data 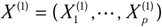, where 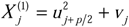, *j* = 1,…, *p*/2, *u_j_* ~ *unif*(−2, 2)and *V_j_* ~ *N*(−4/3, 1/4); 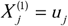 *j* = *p*/2+1,…, *p*, *p* = 100.

We compared JDINAC with several existing state-of-the-art methods under the aforementioned 4 scenarios in differential network analysis and classification.

#### Differential network

We compare the performance of JDINAC in terms of differential network estimation with DiffCorr (Fukushima, 2013), DEDN (Zhao *et al.*, 2014) and cross-product penalized logistic regression (cPLR). The cPLR is defined as

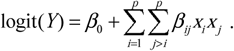

The *L*_1_ penalty function was used to optimize the parameters, which is the same for JDINAC. Parameters *β_ij_* ≠ 0 indicate differential dependency patterns between two groups.

True discovery rate (*TDR*; Precision), true positive rate (*TPR*; Recall), and true negative rate (*TNR*) are used to evaluate the performance of different methods. *TDR*, *TPR* and *TNR* are defined as follows,

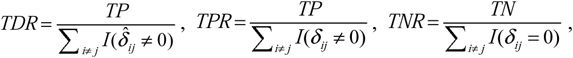

where *TP* and *TN* are the numbers of true positives and true negatives respectively, which are defined as 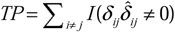, 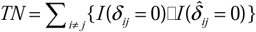 respectively. (*δ_ij_*)_*p*×*p*_ is the differential adjacency matrix, *δ_ij_* ≠ 0 indicate the pair (*x_i_*, *x_j_*) are differential dependency between two groups; 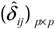 is the estimated differential adjacency matrix.

#### Classification

We compare the classification performance of JDINAC with Random Forest, Naive Bayes, cPLR and original penalized logistic regression (oPLR).The oPLR is defined as

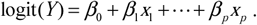

Similarly, the *L*_1_ penalty function was used to optimize the parameters for high-dimensional data. Both cPLR and oPLR are Lasso based methods.

Receiver operating characteristic (ROC) curve and classification error are used to assess the accuracy of 4 methods.

### 2.2 Application

Breast Invasive Carcinoma (BRCA) is the most common type of breast cancer. This subtype of breast cancer is able to spread to other parts of the body through the lymphatic system and bloodstream, which makes BRCA potentially a highly lethal killer. Most of the genome-wide studies for BRCA focus on identifying differentially expressed genes. However, BRCA is largely determined by a number of genes that interact in a complex network, rather than a single gene perturbed (gene mutation, expression, and methylation etc.). A key but inadequately addressed issue is how to identify the underlying molecular interaction mechanisms. The TCGA BRCA study include 1098 patients, along with their matched mRNA, copy number, methylation and microRNA data. The RNASeq Version 2 expression data and clinical data were downloaded from TCGA through TCGA-Assembler (Zhu *et al.*, 2014). In this study, we select 114 patients who have both tumor and matched normal samples as our training subjects. The proposed method was applied to identify differential patterns of network activation between the tumor group and the control group. We focus on the 397 genes listed in the cancer pathway (hsa05200) of KEGG as our candidate gene sets. After filtering those genes which include more than 30% of zero gene expression values in the training data, we have373 genes as our final candidate genes. To evaluate the performances of classification, we randomly choose 50 of 114 individuals in each group as our test data set. More detailed data description and processing is provided in Supplementary.

## 3 Results

### 3.1 Simulation

We calculate the TDR, TPR and TNR of identifying the differential network that corresponds to a given threshold by varying thresholds from 1 to 20 (number of random split was set to be 20 in the Step 4). We average those measures over 100 datasets in each of the 4scenarios.

Table 1 presents the TPR, TNR and TDR of the JDINAC, DiffCorr, DEDN and cPLR under different scenarios. It shows that JDINAC significantly outperforms all the other 3 methods. Although DiffCorr was set to control the false discovery rate (FDR) less than 0.1, the FDR tended to be significantly inflated. In particular, JDINAC performs quite well in scenario 4. The TDR, TPR and TNR of JDINAC are close to 1, but the TDR and TPR of the other 3 methods are close to 0. It indicates that JDINAC can indeed capture the perturbation of nonlinear dependence in the network.

**Table 1.** The TPR, TNR and TDR of different methods. (average of 100 replications, %), The best performance is highlighted in bold.

| Scenario | JDINAC<sup>a</sup> |  |  | DiffCorr<sup>b</sup> |  |  | DEDN |  |  | cPLR |  |  |
| --- | --- | --- | --- | --- | --- | --- | --- | --- | --- | --- | --- | --- |
|  | TDR | TPR | TNR | TDR | TPR | TNR | TDR | TPR | TNR | TDR | TPR | TNR |
| 1 | **93.7** | 81.6 | **99.9** | 81.3 | **100** | 99.8 | 33.5 | 96.7 | 99 | 19.8 | 64.9 | 97.3 |
| 2 | **95.6** | 74.5 | **99.9** | 85 | **100** | 99.7 | 16.5 | 89.1 | 94.3 | 25.6 | 49.8 | 97.1 |
| 3 | **88.3** | **69.5** | **99.3** | 7.5 | 0.2 | 99.8 | 2.1 | 10.1 | 81.6 | 53.6 | 23.6 | 98.1 |
| 4 | **99.9** | **99.8** | **100** | 3.8 | 0.4 | 99.9 | 5 | 0.2 | **100** | 0.7 | 0.3 | 99.8 |

a. Pair $(x_i, x_j)$ was taken as differential edge in the network for JDINAC, when the differential dependency weight $w_{ij} \geq 4$ .
b. Set to control the false discovery rate equal to 0.1.

Figure 3 illustrates the precision-recall curve of JDINAC under different scenarios. The JDINAC has high precision-recall curve in all scenarios.

**Fig 3.**
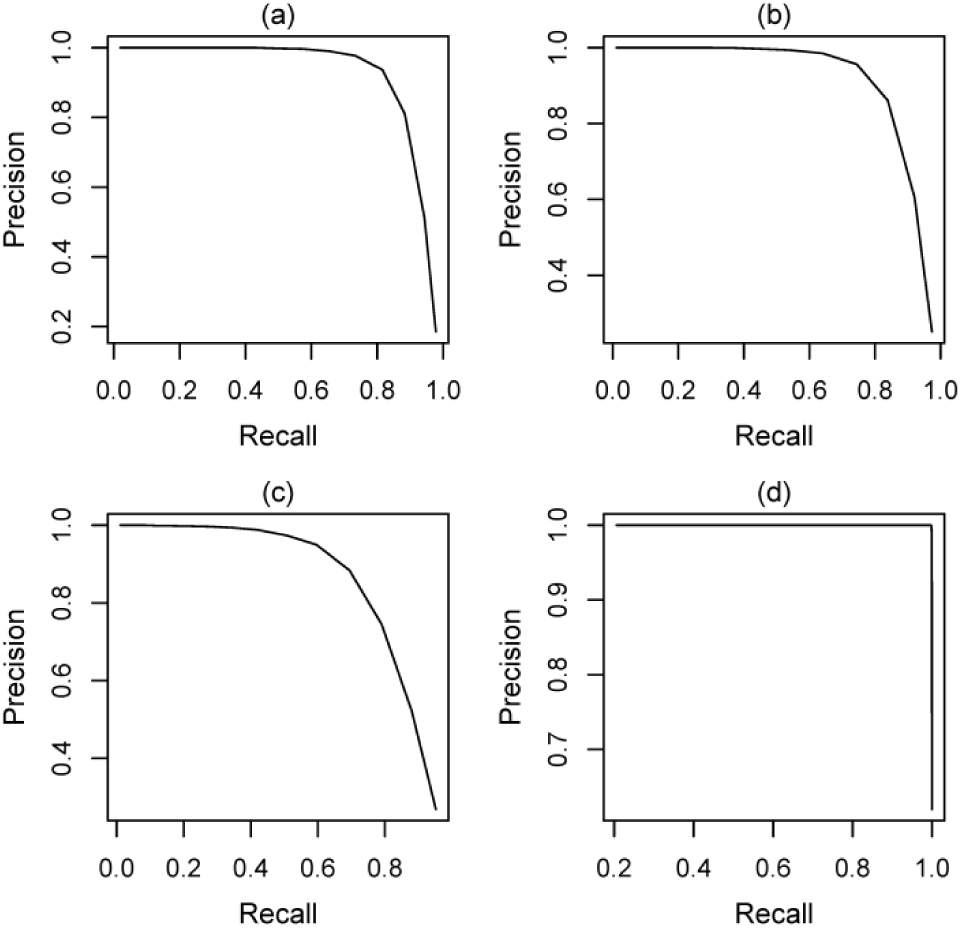
Precision-recall curve for JDINAC for differential network analysis under scenario 1 (a), scenario 2 (b), scenario 3 (c), scenario 4 (d). The differential dependency weights *w_ij_* were used as the differential adjacency matrix, 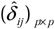 = *I*(*w_ij_* ≥ *t*), *t* = 1,…, 20.

The average ROC curves over 100 replications for the classification using 5 methods under different scenarios (Figure 4) show that JDINAC performs the best among the 5 methods. The fractions of votes were used as the continuous predictions for RF models. After getting the continuous prediction, we used 0.5 as the cutoff of prediction to obtain the classification errors (Table 2). JDINAC is much more accurate than other methods.

**Fig 4.**
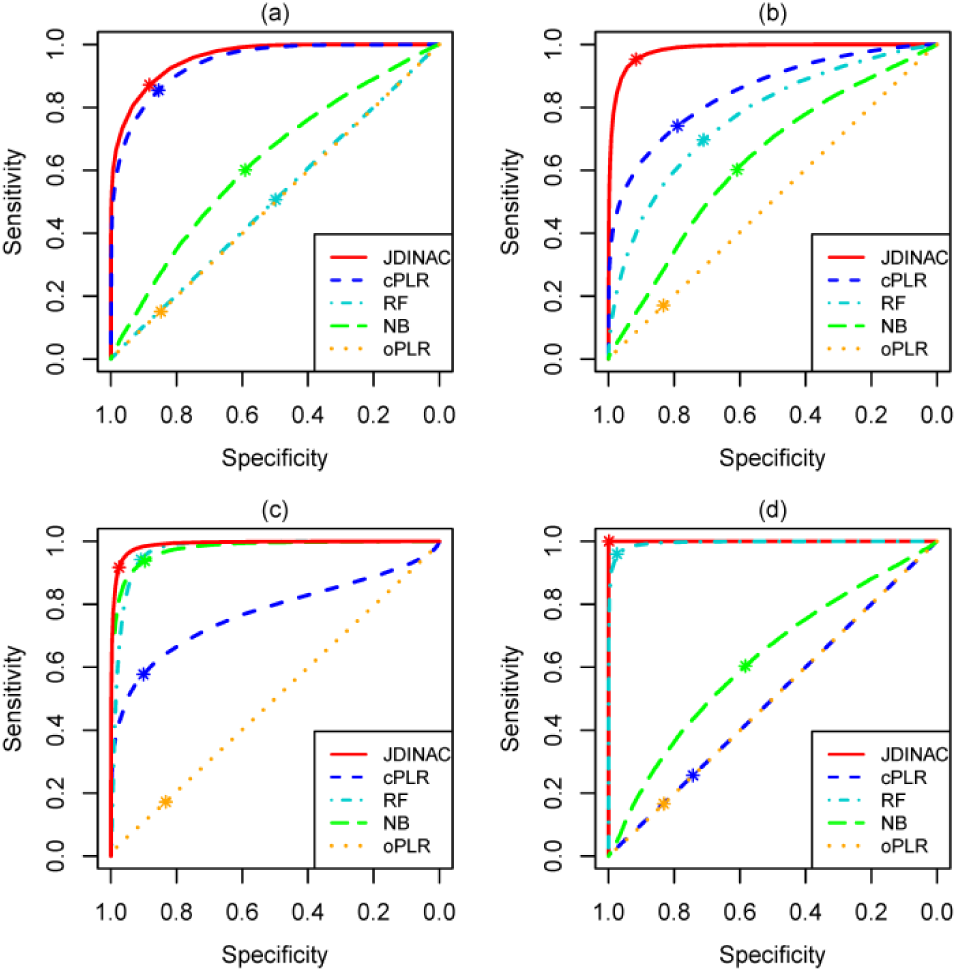
ROC curves of 5 methods for the classification under different scenarios. The asterisk indicates the location where the cutoff of prediction was set to 0.5.

**Table 2.**
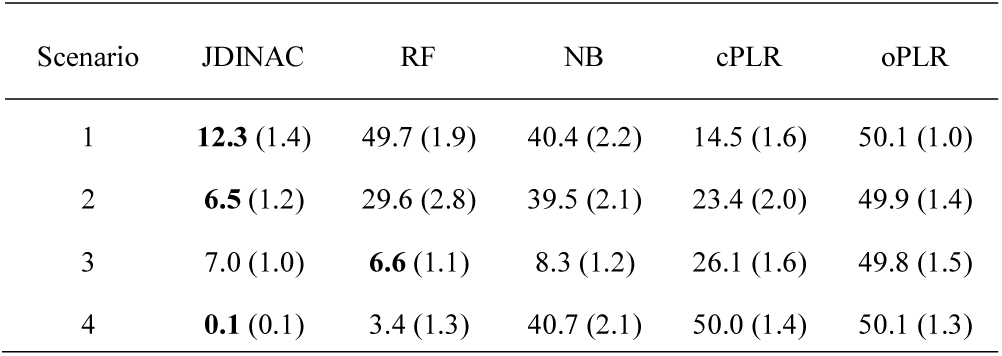
Average classification errors (%). Standard errors are in the parentheses. The best performance is highlighted as bold.

| Scenario | JDINAC | RF | NB | cPLR | oPLR |
| --- | --- | --- | --- | --- | --- |
| 1 | **12.3** (1.4) | 49.7 (1.9) | 40.4 (2.2) | 14.5 (1.6) | 50.1 (1.0) |
| 2 | **6.5** (1.2) | 29.6 (2.8) | 39.5 (2.1) | 23.4 (2.0) | 49.9 (1.4) |
| 3 | 7.0 (1.0) | **6.6** (1.1) | 8.3 (1.2) | 26.1 (1.6) | 49.8 (1.5) |
| 4 | **0.1** (0.1) | 3.4 (1.3) | 40.7 (2.1) | 50.0 (1.4) | 50.1 (1.3) |

### 3.2 Application

Figure 5 depicts the differential network estimated by JDINAC, cPLR, RF, NB and oPLR. Only genes connected with at least one other gene were included in the figure. The top 10 differential dependency pairs identified by JDINAC ordered by weight are shown in Table 3. Figure 6 presents the Venn diagram for the edges in the differential networks identified by different methods DiffCorr, cPLR, and DEDN. There are few overlaps of predicted differential interactions (edges in the network) among these methods. Thus, JDINAC may identify complementary information to the existing methods. The overlapped edges between JDINAC and DiffCorr, JDINAC and cPLR, and DiffCorr and cPLRare shown in Supplementary Table S1.

**Fig 5.**
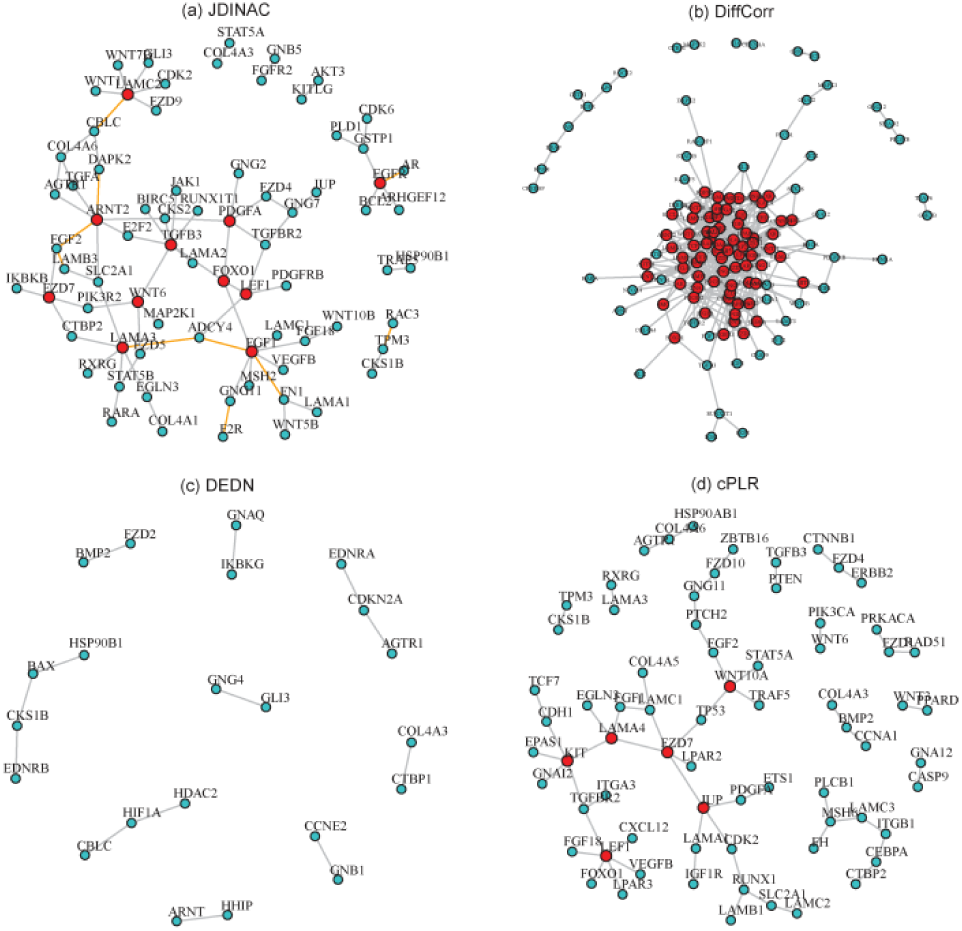
The differential network of cancer pathway between BRCA tumor samples and controls. An edge presented in the differential network means the dependency of corresponding pair genes is different between two condition-specific groups. The red nodes stand for hub genes. (a) Differential network estimated by JDINAC; The orange edges indicate the top 10 differential dependency pairs. (b) Differential network estimated by DiffCorr; (c) Differential network estimated by DEDN; (d) Differential network estimated by cPLR.

**Table 3.** Top 10 differential dependency pairs identified by JDINAC.

| Gene 1 | Gene 2 | $w_{1,2}$ | Gene 1 | Gene 2 | $w_{1,2}$ |
| --- | --- | --- | --- | --- | --- |
| *GNG11* | *F2R (PAR1)* | 18 | *LAMA3* | *ADCY4* | 12 |
| *FN1* | *FGF1* | 17 | *EGFR* | *AR* | 10 |
| *LAMB3* | *FGF2* | 17 | *DAPK2* | *ARNT2* | 10 |
| *TPM3* | *RAC3* | 17 | *FGF2* | *ARNT2* | 10 |
| *FGF1* | *ADCY4* | 13 | *LAMC2* | *CBLC* | 10 |

**Fig 6.**
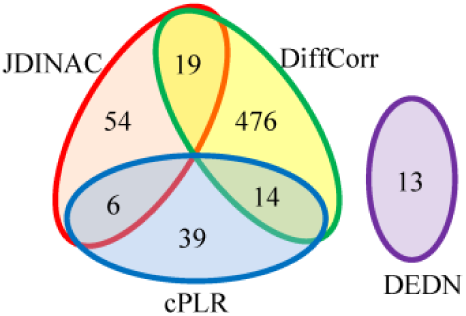
Summary of the number of edges in the differential networks for the 4 methods.

No gold standard is available for evaluating differential network analysis in the real dataset since the true underlying dependence relationships are unknown. Therefore, we can no longer study the performance of JDINAC in terms of *TDR*, *TPR* and *TNR* in the real data analysis. Supplementary Table S2 presents the hub genes and the corresponding number of neighbor genes identified by JDINAC. The hub genes are the ones that have at least 3 neighbor genes in the differential networks.

Next, we study the classification performances of methods JDINAC, RF, NB, cPLR and oPLR.The classification errors are shown in Table 4. The classification accuracy of JDINAC is the same with oPLR that uses single genes as features, but better than RF, NB and cPLR, all of which use the pair of genes for the classification The low error rate of JDINAC implies that the identified differential network could be biological meaningful to distinguish the disease state with the normal one.

**Table 4.** Classification errors on application test data set (%).

| Dataset | JDINAC | RF | NB | cPLR | oPLR |
| --- | --- | --- | --- | --- | --- |
| BRCA | **1** | 19 | 2 | 17 | **1** |

## 4 Discussion

A complex disease phenotype (e.g. cancer) is rarely a consequence of an abnormality in a single gene, but reflects various pathobiological processes that interact in a network (Barabasi *et al.*, 2011). Network comparison or differential network analysis has become an important means of revealing the underlying mechanism of pathogenesis. The identified differential interaction patterns between two group-specific biological networks can be taken as candidate biomarkers, and have extensive biomedical and clinical applications (Ji *et al,* 2015, 2016; Laenen *et al,* 2013; Yang *et al.*, 2013). Although numerous differential network analysis methods (Fukushima, 2013; Ha *et al,* 2015; Watson, 2006; Yates and Mukhopadhyay, 2013; Zhao *et al.*, 2014) have been proposed, most of the methods rely on marginal or partial correlation to measure the strength of connection between pairs of nodes in a network. They usually cannot capture the nonlinear relationship among genes, which could be ubiquitous in real applications.

We propose a joint kernel density based method, JDINAC, for identifying differential interaction patterns of networks between condition-specific groups and conducting discriminant analysis simultaneously. A multiple splitting and prediction averaging procedure were employed in the algorithm of JDINAC. It can not only make the approach more robust and accurate, but also make more efficient use of limited data (Fan *et al.*, 2016). Moreover, the nonparametric kernel method was used to estimate the joint density, which does not require any conditions on the distribution of the data; this also makes JDINAC more robust and has the ability to capture the nonlinear relationship among genes. Extensive simulations were conducted to assess the performances of differential network analysis and classification accuracy. It indicated that JDINAC has high reliability (Figure 2) and significantly outperforms other state-of-the-art methods, DiffCorr, DEDN and cPLR, especially in scenarios 3 and 4 for the differential network analysis (Table 1). One advantage for JDINAC is that it can achieve classification simultaneously, making it more attractive in practical applications. Figure 3 and table 2 further highlighted that JDINAC is much more accurate in classification than other methods.

JDINAC was applied to BRCA dataset, the differential network was estimated and several hub genes were found. We found there are experimental support for the top ranked pairs by JDINAC. For example, *F2R* (*PAR1*) is a G protein coupled receptor (GPCR) that binds and regulates G-protein. It contributes to tumor progression and metastasis in breast cancer (Shi *et al.*, 2004). Meanwhile, *GNG11* is a G-protein, plays a role in the transmembrane signaling system. It implies that the molecular role of *F2R* in the breast cancer progression and metastasis origins from the altered *F2R-GNG11* interaction. In other cases, dysregulated pairs may not have direct physical interactions, but strong functional associations. The matrix form of fibronectin (*FN1*) is believed to support cell adhesion, tumor growth, and inflammation. Fibroblast growth factors (*FGF1*, *FGF2*) are important factors regulating expression of *FN1* and *LAMB3* (Kashpur *et al.*, 2013; Tang *et al.*, 2007). *RAC3* is a GTPase which is related to the cell growth and the activation of protein kinases. RacGTPase activity and paxillin phosphorylation are elevated in cells from the *TPM3*tropomyosin gene deleted mice(Lees *et al.*, 2013).

*FGF1* and *TGFB3* have the largest number of neighbor genes in the differential networks of BRCA data. *FGF1* plays an important role in a variety of biological processes involved in embryonic development, cell growth and differentiation, morphogenesis, tumor growth and invasion (Zhou *et al.*, 2011). The expression of *FGF1* is dysregulated in breast cancer and contributes to the proliferation of breast cancer cells(Yoshimura *et al.*, 1998; Zhou *et al.*, 2011). Laverty *et al.* (Ghellal *et al.*, 2000) reviewed numerous literatures and reported *TGFB3* is associated with the progression of breast cancer. *PDGFA* is confirmed to be one of the progesterone target genes on breast cancer cells (Soares *et al.*, 2007). *FOXO1* contributes to multiple physiological and pathological processes including cancer, and targeting of *FOXO1* by microRNAs may promote the transformation or maintenance of an oncogenic state in breast cancer cells(Fu and Tindall, 2008; Guttilla and White, 2009). Moreover, *FOXO1* is regulated by *AKT* (Tzivion *et al.*, 2011), and *PDGFA* is the upstream gene of *AKT*. Indeed, we identified an edge between *PDGFA* and *FOXO1* (Figure 4a). Wendt M K *et al.* (2015) demonstrated that *EGFR* is a critical gene in primary breast cancer initiation, growth and dissemination.*FZD7* plays a critical role in cell proliferation in triple negative breast cancer (TNBC) via Wnt signaling pathway and was considered to be a potential therapeutic target for TNBC (Yang *et al.*, 2011). An edge between *FZD7* and *CTBR2* was identified by JDINAC (Figure 4a). Actually, *CTBP2* is a key gene in Wnt pathway. The identified differential network provides new insight into the underlying genetic mechanisms of BRCA, and testable hypothesis for further experimental validations. The differential interaction patterns and hub genes may serve as biomarkers for early diagnosis or drug targets.

Although JDINAC can in principle be applied to genome-wide data sets, such application may be limited due to high computational costs. In this study, we focus on identifying the differential interaction patterns between genes in a given pathway (or a candidate gene set). JDINAC can be directly used in most cases, since more than 95% pathways from KEGG database contain less than 150 genes. Under the scenario when the pathway is too large or in the case of genome-wide study, prior knowledge or screening method can be used to shrink the candidate gene pair numbers before applying JDINAC. Although the proposed JDINAC method was applied to gene network differential network analysis in this paper, it can be used to incorporate other biological networks, such as metabolic network and brain functional connectivity network. It can also be generalized to identify of between pathway interactions.

The freely available JDINAC software is available as R script at https://github.com/jijiadong/JDINAC.

## Acknowledgements

The authors would like to acknowledge TCGA for providing the BRCA data. We would also like to thank the patients for access to their study data.

